# Bulk segregant analysis reveals environment × genotype interactions determining malaria parasite growth

**DOI:** 10.1101/2020.09.12.294736

**Authors:** Sudhir Kumar, Xue Li, Marina McDew-White, Ann Reyes, Elizabeth Delgado, Abeer Sayeed, Meseret T. Haile, Biley A. Abatiyow, Spencer Y. Kennedy, Nelly M. Camargo, Lisa A. Checkley, Katelyn V. Brenneman, Katrina A. Button-Simons, Manoj T. Duraisingh, Ian H. Cheeseman, Stefan H. I. Kappe, François Nosten, Michael T. Ferdig, Ashley M. Vaughan, Tim J. C. Anderson

## Abstract

What genes determine growth and nutrient utilization in asexual blood-stage malaria parasites? Competition experiments between a lab-adapted African parasite (NF54), and a recently isolated Asian parasite (NHP4026) reveal contrasting outcomes in different media: NF54 outcompetes NHP4026 in media containing human serum, while NHP4026 outcompetes NF54 in media containing AlbuMAX, a lipid-rich bovine serum formulation. We conducted parasite genetic crosses and compared genome-wide allele frequency changes in progeny populations cultured in media containing serum or AlbuMAX: this bulk segregant analysis (BSA) reveals three quantitative trait loci (QTL) underlying differential growth. The strongest QTL (chromosome 13) contains *EBA-140*: competition experiments between *EBA-140*-knockout and isogenic wildtype parasites showed fitness reversals in the two media types, validating this locus as the causative gene. These results (i) demonstrate the effectiveness of BSA for dissecting fitness traits in *Plasmodium falciparum*, and (ii) reveal an intimate link between red blood cell invasion and nutrient composition of growth media.

---

Asexual blood stage malaria parasite growth rates are determined by factors including efficiency of red blood cell (RBC) invasion, nutrient acquisition and proliferation within red blood cells. Not surprisingly, genes underlying these processes are promising targets for vaccine and antimalarial development. For example, parasite genes underlying RBC invasion, such as apical membrane antigen (*AMA1*), are the focus of vaccine development efforts^1^. Among existing drugs, artemisinin (ART) - the frontline drug against malaria - is activated by hemoglobin digestion^2, 3, 4^, while chloroquine (CQ) interferes with heme polymerization into non-toxic hemozoin^5^, and antifolate drugs (pyrimethamine and sulfadoxine) are competitive inhibitors that interrupt the folate biosynthesis pathway^6^. The mutations conferring resistance to these drugs are also involved with parasite nutrition transport/metabolism pathways. For example, resistance to ART, is mediated by mutations in *kelch13*, which is required for hemoglobin endocytosis^2^. Mutations in the CQ resistance transporter (*PfCRT*), which normally functions as a transport channel for ions and peptides mediate resistance to a variety of drugs including CQ and piperaquine^5^. Mutations in dihydrofolate reductase and dihydropteroate synthase, components of the folate synthesis pathway confer resistance to pyrimethamine and sulfadoxine^6^. Given the importance of RBC invasion and nutrient acquisition/metabolism, effective methods for locating genes and pathways involved in these processes are urgently needed.

Exploiting natural variation in parasite nutrient acquisition and metabolism pathways provides one promising approach. Nguitragool *et al*. used a human malaria parasite *Plasmodium falciparum* genetic cross conducted in chimpanzee hosts to identify an important channel (plasmodial surface anion channel, PSAC) involved in ion transport^7^. Similarly, Wang *et al*. used a comparable linkage mapping approach to investigate the ability of parasites to utilize exogenous folate^8^, as this is important for determining the success of drugs that target the parasite folate synthesis pathway. These linkage analysis studies demonstrate how differences in metabolism or nutrient acquisition between parasites can be effectively exploited to better understand the genetic underpinning of metabolic pathways and transport systems, which in turn can highlight potential targets for intervention. However, traditional linkage mapping, requiring phenotyping and sequencing of individual cloned progeny, is laborious and does not access the full power of independent recombinants available in uncloned populations.

Our central aim was to evaluate the efficacy of genetic crosses and a rapid linkage mapping method - bulk segregant analysis (BSA) - for understanding the genetic basis of nutrition-related phenotypes in *P. falciparum* using differential growth and fitness in serum- or AlbuMAX-based *in vitro* culture as a test system. BSA provides a simple and fast approach to identify loci that contribute to complex traits^9^ that are typically identified using traditional QTL mapping. Using sequencing of pooled progeny populations, BSA measures changes in allele frequency following distinct selections (here comparing asexual blood stage growth in serum or AlbuMAX). BSA, also referred to as linkage group selection (LGS), has been extensively used with genetic crosses of rodent malaria parasites to map genes determining blood stage multiplication rate, virulence and immunity in *Plasmodium yoelii*^10, 11^, as well as mutations conferring ART resistance and strain-specific immunity in *Plasmodium chabaudi*^12^. Carrying out genetic crosses in *Anopheles* mosquitoes and human hepatocyte-chimeric mice (FRG huHep mice)^13^ now allows us to routinely generate large pools of recombinant *P. falciparum* progeny without the need for chimpanzee hosts. We have previously generated four independent *P. falciparum* genetic crosses with large numbers of unique recombinant progeny using this approach (^14^, reviewed in^15^).

Continuous *in vitro* culture of asexual blood stages of the malaria parasite *P. falciparum* typically requires human RBCs, buffered RPMI 1640 medium and human serum, with a low oxygen atmosphere at 37°C^16^. RPMI 1640 medium is the main resource for sugar (glucose), salts, essential amino acids and multiple vitamins^17^. Human hemoglobin can supply amino acids other than isoleucine^18^, while human serum provides most of the other nutrients needed for parasite growth, such as inorganic and organic cations. Lipid-enriched bovine albumin (AlbuMAX) is the most widely used human serum substitute and extensively used for blood stage parasite culture. Human serum typically contains more phospholipid and cholesterol, and less fatty acid than AlbuMAX^19^. AlbuMAX has several advantages over human serum for culture of *P. falciparum* due to its low cost, compatibility with any blood type, and lower batch-to-batch variation. AlbuMAX-supplemented culture medium has made significant contributions to malaria research, facilitating *in vitro* drug sensitivity assays for screening and monitoring of antimalarial drugs, parasite growth competition assays to measure fitness costs, and research on parasite molecular biology and immunology. However, several studies have found parasite growth rate differences between AlbuMAX- and human serum-based cultures and these can impact drug susceptibility profiling. For example, AlbuMAX supported parasite growth less well than human serum for clinical isolates from Cameroonian patients^20^, and for long-term lab culture adapted parasites^21^. Furthermore, the 50% inhibitory concentrations (IC50s) of multiple antimalarial drugs obtained with AlbuMAX, including CQ, amodiaquine, quinine and ART, were almost twice the corresponding values obtained with human non-immune serum^22^.

This study is founded on the observation that parasites show profound differences in competitive growth in serum and AlbuMAX: NF54 outcompetes NHP4026, a parasite clone from Thailand^23^, in media containing human serum, while NHP4026 outcompetes NF54 in media containing AlbuMAX. We therefore generated replicate genetic crosses between these two parasites, and measured changes of allele frequency in independent progeny populations during parallel asexual blood stage growth in serum and AlbuMAX. This revealed three repeatable quantitative trait loci (QTL) regions, on chromosomes 2, 13 and 14, that were associated with large differences in parasite growth rates in the two media types. Finally, using knockout parasites, we showed that an RBC invasion gene, erythrocyte binding antigen-140 (*EBA-140*), causes the largest QTL for this environment **×** genotype interaction.

## Results

### The outcome of parasite competition is reversed in different media

NF54 is a parasite of Africa origin^24^ that has been maintained in the lab for decades, in a variety of media formulations containing both serum and AlbuMAX. NHP4026 was cloned from a patient visiting the Shoklo Malaria Research Unit (SMRU) clinic on the Thailand-Myanmar border in 2007 and has been cultured in the lab in media containing AlbuMAX for ~80 days.

We measured the competitive fitness of NF54 and NHP4026 blood stage parasites in media containing human serum, AlbuMAX, or a combination of these media types (Fig. 1). NF54 outcompeted NHP4026 in media containing human serum, while NHP4026 outcompeted NF54 in media containing AlbuMAX, and the two parasites showed comparable fitness in media containing both human serum and AlbuMAX (Fig. 1A). The fitness differences observed in human serum and AlbuMAX were large. NF54 replaced NHP4026 in five asexual life cycles in media containing serum: this resulted from a selective advantage (*s*) of 0.51±0.15 (1× standard error) (replicate 1) and 0.62±0.07 (replicate 2) per 48 h asexual cycle (Fig. 1B). In contrast, NHP4026 replaces NF54 in eight asexual life cycles in AlbuMAX, and the relative fitness is reversed (*s* = −0.13±0.05 for replicate 1 and −0.14±0.04 for replicate 2) (Fig. 1B). The differences in fitness observed between the two media types were highly significant (*p* = 7.37E-10).

**Fig 1.**
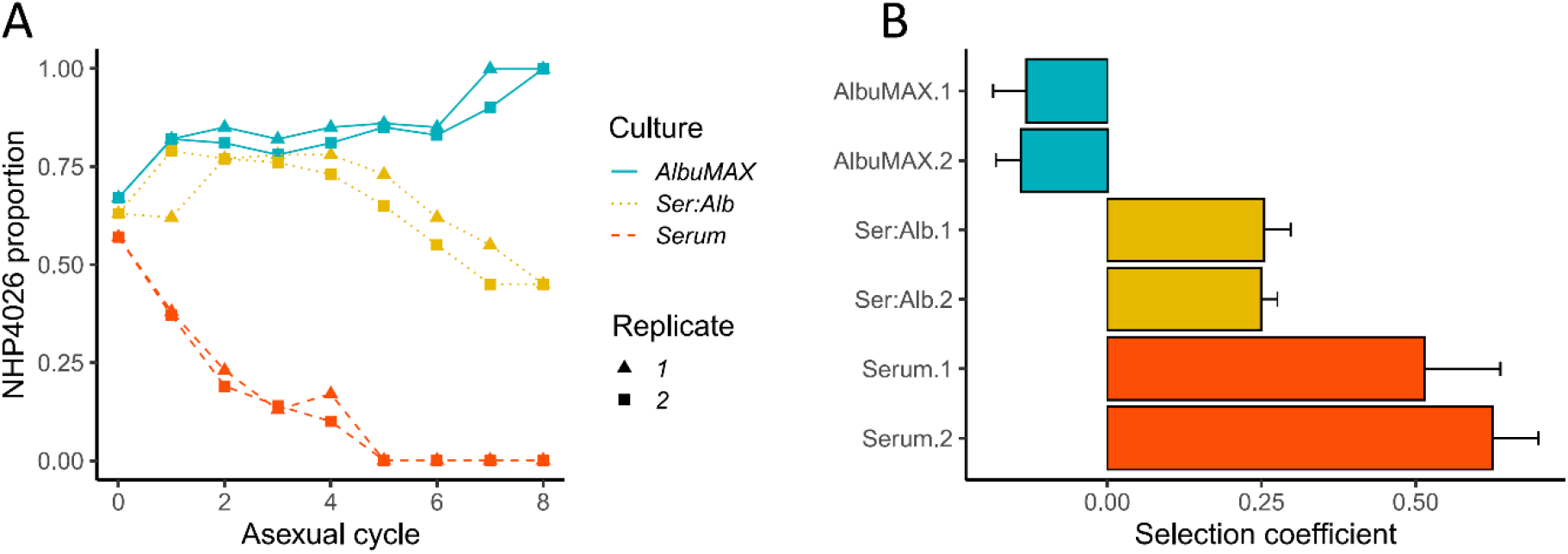
Outcome of competition between NF54 and NHP4026 under different culture conditions. (A), Proportion of NHP4026 against parasite asexual cycles. (B), selection coefficients (*s*) of NHP4026 with 1× standard error. Positive values of *s* indicate a disadvantage for alleles inherited from NHP4026. Culture conditions: AlbuMAX, culture media only contain AlbuMAX not serum; Ser:Alb, culture media contain both serum and AlbuMAX at 50:50 ratio; Serum, culture media contain only serum not AlbuMAX.

### Genetic crosses and composition of segregant pools

We generated crosses between NF54 and NHP4026 (Fig. 2A). There are a total of 13,195 single nucleotide polymorphisms (SNPs, Supplementary Table 1) differentiating the two parental parasites within the 21 Mb core genome (defined in^25^). We used *Anopheles stephensi* mosquitoes and FRG huHep mice^13,26^ to generate three independent recombinant pools containing an estimated 2800 unique recombinants per pool (see Methods for details). Whole genome sequencing (WGS) data of cloned progeny from a mixture of all pools revealed low numbers of selfed progeny (2/55) in this cross with very little redundancy among recombinants (Supplementary Table 2) as observed in previous crosses^12^. We cultured each recombinant pool with O-positive non-immune human serum or AlbuMAX for 34 days in parallel, with two technical replicates in each media type for each of the three biological replicates. We collected samples for BSA every four days and used WGS to analyze all segregant pools to high (191 ± 40 read depth) genome coverage (Fig. 2B).

**Fig 2.**
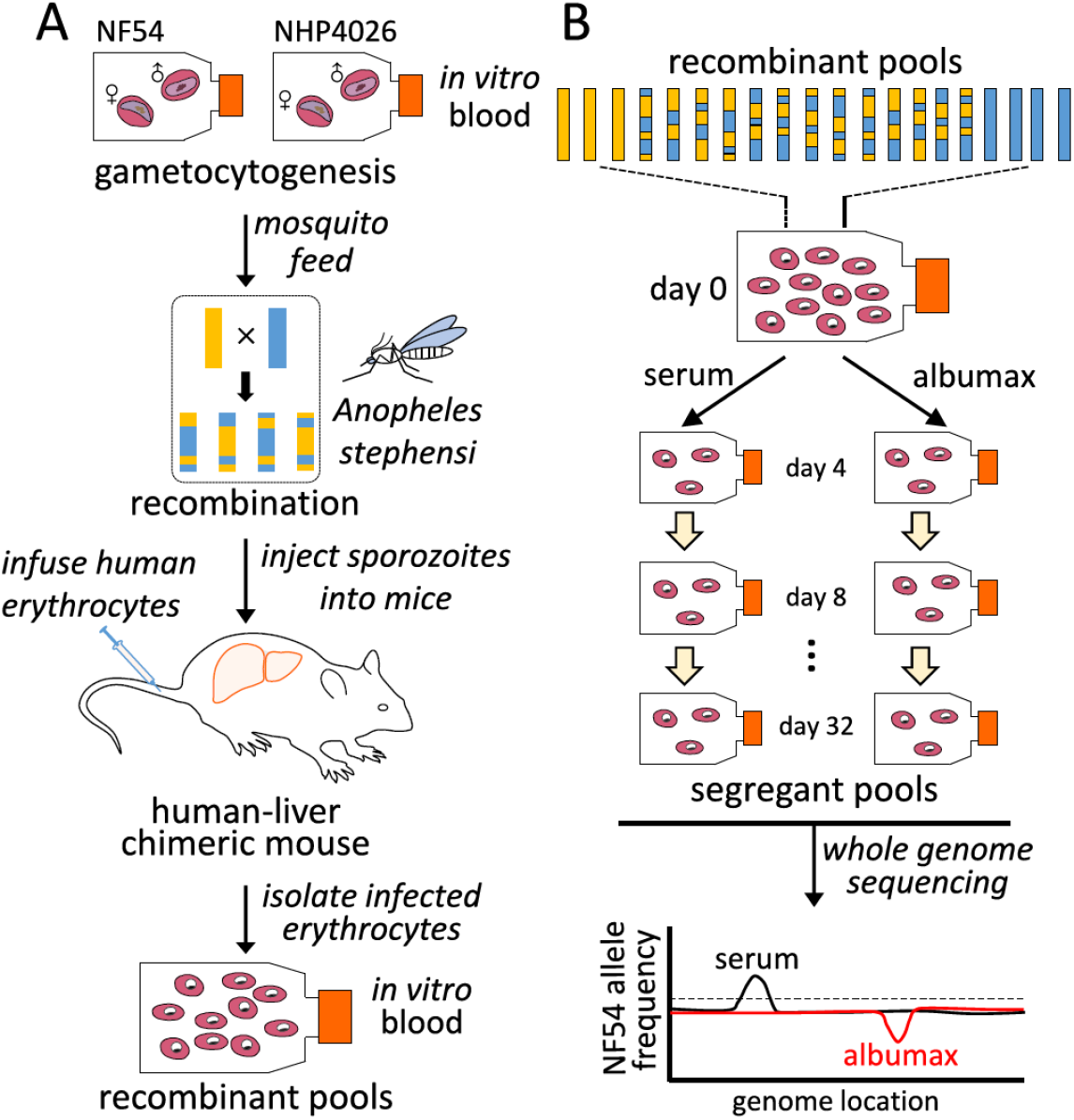
Mapping parasite fitness under different culture conditions. **(A)**, Recombinant progeny pool generation. Genetic crosses are generated using female *Anopheles stephensi* mosquitoes and FRG huHep mice as described by Vaughan et al (*13*). Recombination of parasite genomes occurs during meiosis in the mosquito midgut. Recombinant pools are collected from infected mice and maintained through *in vitro* blood cultures. (**B)**, Bulk segregant analysis. Pools of progeny are cultured in parallel with serum or AlbuMAX and samples are collected every 4 days. Each segregant pool is then whole-genome sequenced and genotyped in bulk. Differences in allele frequency among different groups are used to identify QTLs.

**Table 1.** Top candidate genes located inside of the QTL regions.

| **QTL ID** | **QTL<sup>a</sup>** | **95% CI<sup>b</sup>** | **Gene ID** | **Gene name** | **Gene annotation** |
| --- | --- | --- | --- | --- | --- |
| *QTLs for differential growth in serum and AlbuMAX* |  |  |  |  |  |
| chr.2 | serum vs<br>AlbuMAX | chr2: 0 – 220,805 | PF3D7_0204500 | *AST* | aspartate transaminase |
| chr.13 | serum vs<br>AlbuMAX | chr13: 0 – 162,844 | PF3D7_1301600 | *EBA-140* | erythrocyte binding antigen-140 |
| chr.14.1 | serum vs<br>AlbuMAX | chr14: 629,528-812,881 | PF3D7_1417300 | *ATG4* | cysteine protease ATG4 |
| *QTLs revealed by allele frequency skews in both serum and AlbuMAX* |  |  |  |  |  |
| chr.7 | serum and<br>AlbuMAX | chr7:340,864 - 476,223 | PF3D7_0709000 | *CRT* | chloroquine resistance transporter |
| chr.12 | serum and<br>AlbuMAX | chr12:1,141,368 - 1,282,609 | PF3D7_1229100 | *MRP2* | multidrug resistance-associated<br>protein 2 |
| chr.14.2 | serum and<br>AlbuMAX | chr14:2,356,428 - 2,485,473 | PF3D7_1460900 | *ARPS10* | apicoplast ribosomal protein S10 |

a, serum vs AlbuMAX QTLs were identified by comparing differential growth in human serum and AlbuMAX; serum/AlbuMAX QTLs were detected in both Serum and AlbuMAX.
b, 95% confidence interval (CI) using Li's method <sup>59</sup>.

### Bulk segregant analysis identifies QTLs for differential growth in human serum and AlbuMAX

By comparing allele frequency changes in human serum and AlbuMAX culture over 34 days (Fig. 3), we detected three QTLs (Table 1, Fig. 4, Supplementary Fig. 1) situated at the beginning of chromosome 2 (named QTL chr.2), the beginning of chromosome 13 (QTL chr.13) and the first half of chromosome 14 (QTL chr.14.1). For each detected QTL (G’ > 20), we calculated 95% confidence intervals to narrow down the size of the genome region and thus the list of genes that could be driving selection (Fig. 5). The list of genes inside the QTL regions is summarized in Supplementary Table 3. We prioritized genes within these genome regions by the following criteria: i) evidence of gene expression in blood stages; ii) gene annotations and related metabolic pathways; iii) inspection of SNPs and indels that differentiated the two parents (Supplementary Table 4 & 5). The strongest candidate genes driving these QTLs are listed in Table 1.

**Fig 3.**
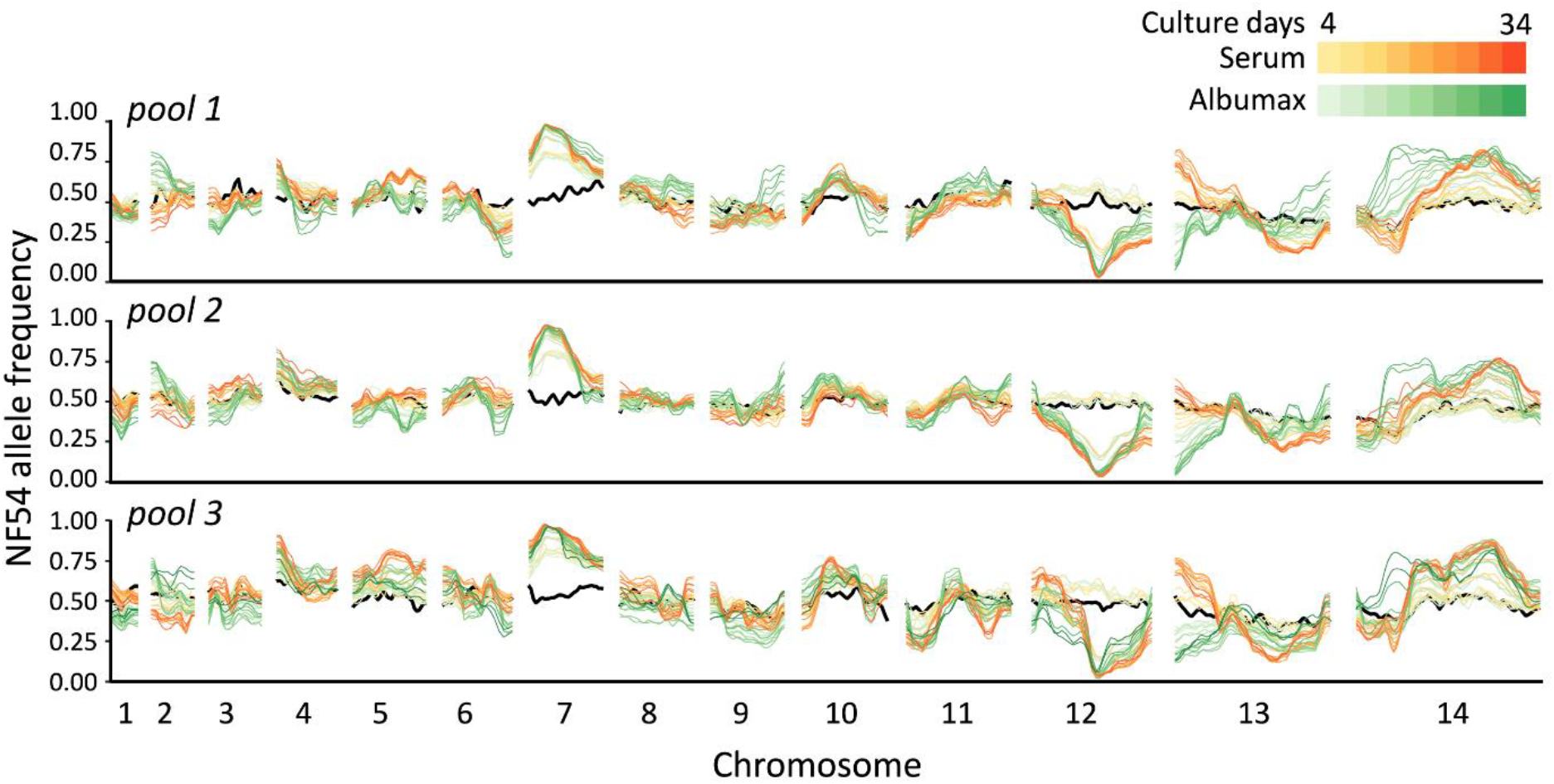
Change in frequency across the genome in different culture conditions. The black lines show allele frequencies from the initial recombinant pools, while red and green lines indicate allele frequency changes during serum (red) or AlbuMAX (green) cultures. The three plots show allele frequencies from different recombinant pools (biological replicates). The lines with same color in each panel show values for the two technical replicates.

**Fig 4.**
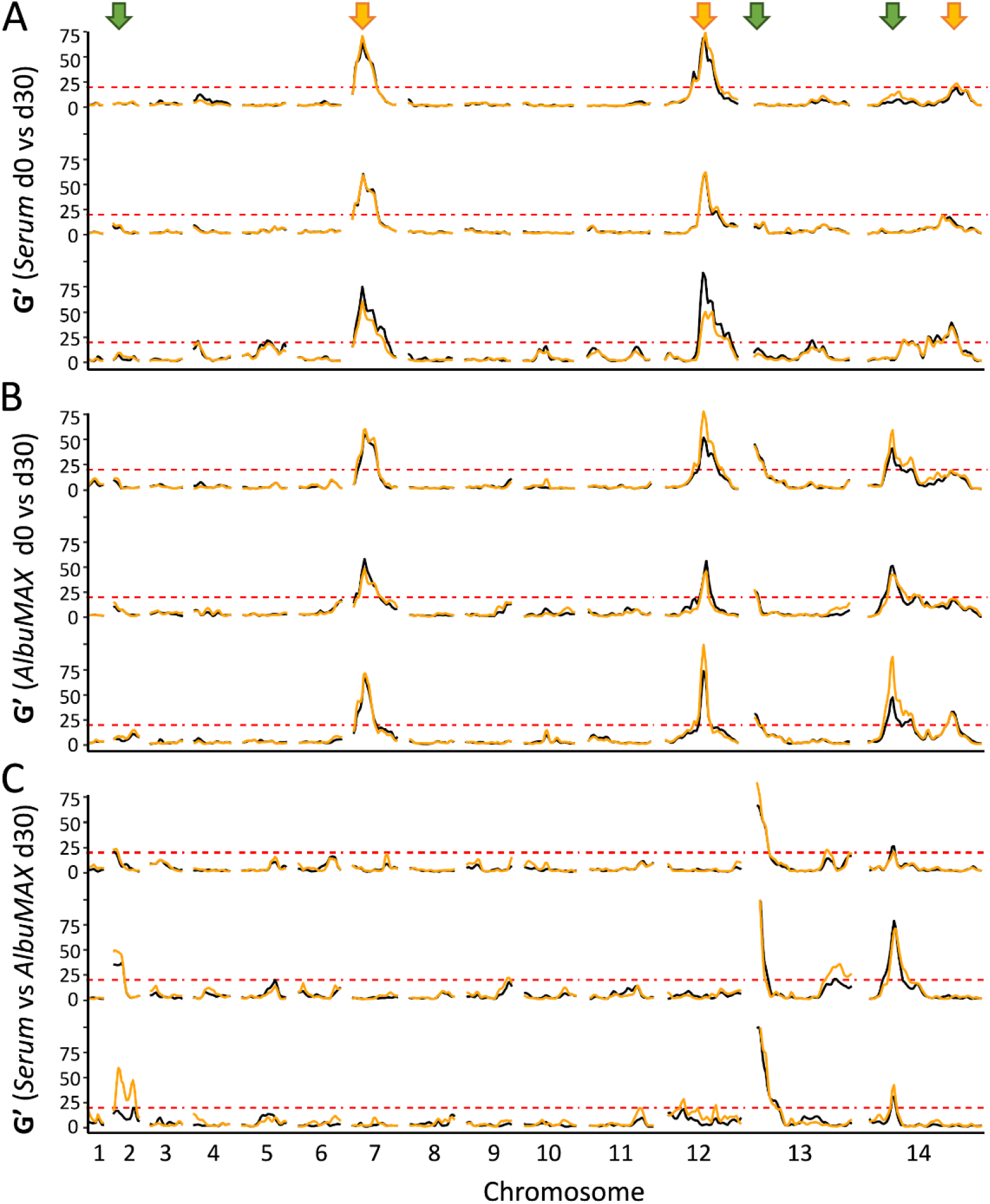
QTLs defined with G’ approach.

The top and middle panels show QTLs detected through comparing allele frequencies from the initial recombinant pools and pools after 30 days of serum (top) or AlbuMAX (middle) culture. Orange arrows (top) mark the position of QTLs. The bottom panel indicates QTLs at day 30 between serum and AlbuMAX cultures. Green arrows(top) mark the position of QTLs. There are three plots in each panel, which are from different recombinant pools (biological replicates). Orange and black lines are technical replicates in each experiment. We used a threshold (G’ > 20) to determine significant QTLs.

**Fig 5.**
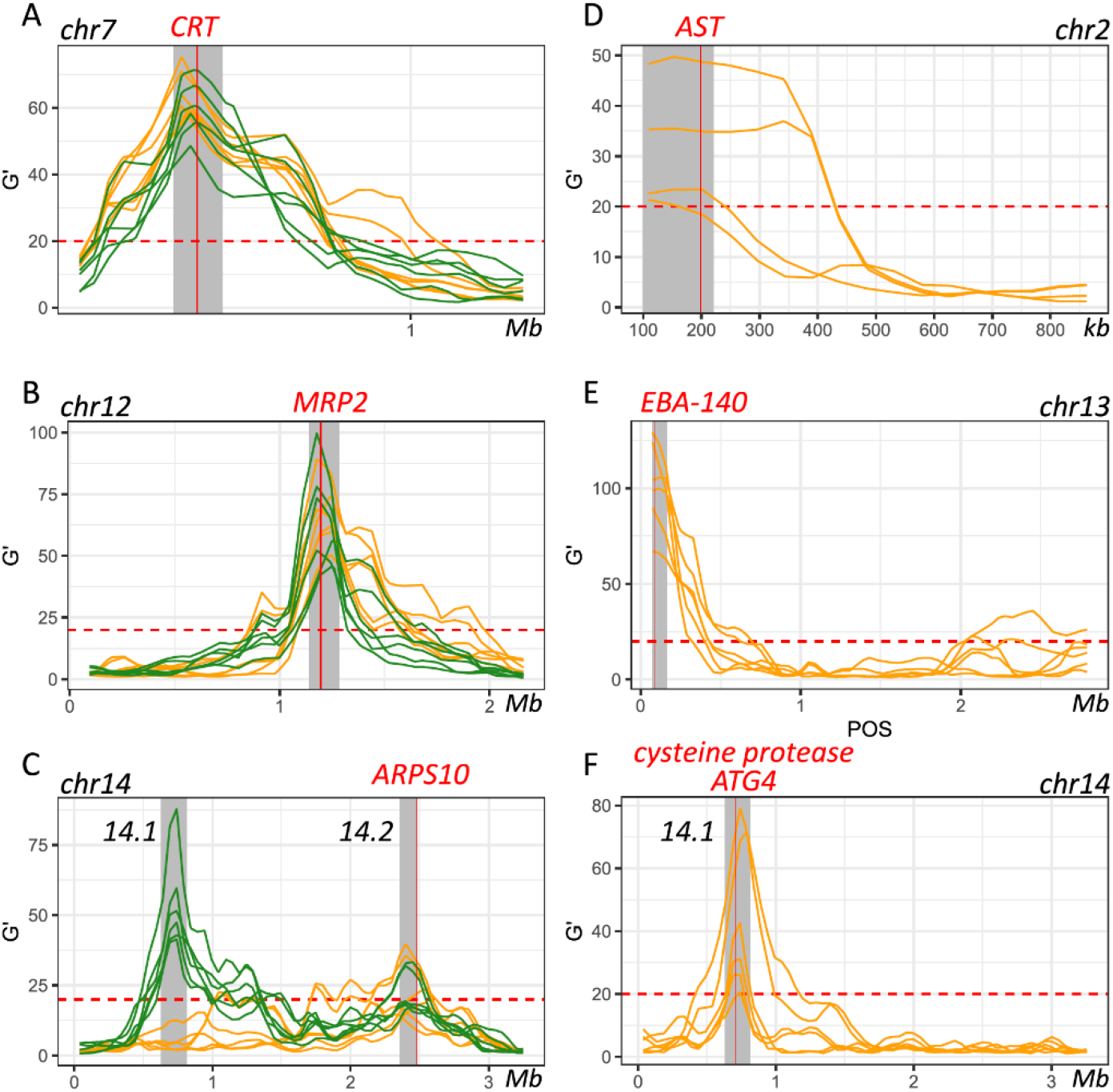
Genes inside QTL regions. (**A**, **B** & **C)**, QTLs detected through comparing allele frequencies from the initial recombinant pools and pools after 30 days of serum (orange) or AlbuMAX (green) culture. (**D**, **E** & **F)**, QTLs by comparing serum and AlbuMAX after 30 days of culture. Each line is one comparison. Grey shadows indicate boundaries of the merged 95% confidential intervals (CIs) of all the QTLs.

The top and middle panels show QTLs detected through comparing allele frequencies from the initial recombinant pools and pools after 30 days of serum (top) or AlbuMAX (middle) culture. Orange arrows (top) mark the position of QTLs. The bottom panel indicates QTLs at day 30 between serum and AlbuMAX cultures. Green arrows (top) mark the position of QTLs. There are three plots in each panel, which are from different recombinant pools (biological replicates). Orange and black lines are technical replicates in each experiment. We used a threshold (G’ > 20) to determine significant QTLs.

For the chr.2 QTL, the NF54 allele frequency increased over time in AlbuMAX and decreased in serum; while the opposite pattern was observed for chr.13 QTL. The patterns of divergent selection observed were consistent across all biological and technical replicates (Supplementary Fig. 2, Supplementary Table 6). For NF54, we observed *s* = 0.04±0.01 in serum and *s* = −0.06±0.01 in AlbuMAX for the chr.2 QTL region; *s* = −0.08±0.02 in serum and *s* = 0.15±0.01 in AlbuMAX for chr.13. These two QTL regions are located at the beginning of the chromosome. It was difficult to define QTL boundaries to a narrow confidence interval due to limited recombination and difficulties of sequence alignment in these regions. For the chr.2 QTL, we inspected genes located from the beginning of chr.2 to 220 kb (chr.2 has a length of 947 kb) (Fig. 5). This region contained 53 genes, 14 are not expressed in blood stage parasites and were excluded, and 13 of the remaining 39 were considered high priority candidates. Among these, the aspartate transaminase gene (*AST*, PF3D7_0204500, also known as aspartate aminotransferase, *AspAT*), is known to be a critical enzyme for amino acid metabolism^27^. There were no non-synonymous mutations in the coding region in the *AST* alleles from the two parents, but we found multiple differences within the 5’ UTR and gene expression regulatory regions, including two SNPs (c.-18C>T [18 bp upstream the coding region] and c.-29A>C) and three microsatellites (Supplementary Table 4).

The chr.13 QTL contained 33 genes (23 expressed in blood stage, 8 high priority candidates) and spanned 163 kb at the beginning of chr.13. Among the genes in this QTL was erythrocyte binding antigen-140 (*EBA-140*, PF3D7_1301600). EBA-140 mediates *P. falciparum* RBC invasion by binding to the RBC receptor glycophorin C and initiating merozoite entry^28^ and expression levels are variable in parasites from different clinical isolates^29^. Interestingly there is a single non-synonymous mutation (Leu112Phe) between NF54 and NHP4026. This SNP is located before the first Duffy-binding-like (DBL) domain^30^ and is common in malaria populations (Supplementary Fig. 3). The *EBA-140* allele from NHP4026 was preferentially selected for in AlbuMAX (Supplementary Fig. 2).

In the chr.14.1 QTL region, the NF54 allele frequency did not change over time during growth in serum but increased significantly during growth in AlbuMAX (Fig. 3). The selection coefficient *s* = 0.02±0.02 in serum and *s* = −0.10±0.03 in AlbuMAX (Supplementary Fig. 2). This QTL, located in the first half of chr. 14 (630 kb – 813 kb, Supplementary Table 5), spanned 183 kb and contained 38 genes. Of these genes, 37 are expressed in blood stages, and 11 are high priority candidates. Interestingly, NHP4026 carried a single amino acid deletion (Asn226del) and three non-synonymous mutations (Ser329Pro, Asn503Lys and Val556Ile) in the cysteine protease *autophagy-related 4* gene (*ATG4*, PF3D7_1417300) within this region.

### Functional validation of EBA-140 as the causative gene within the chr.13 QTL

We chose the strongest QTL on chr. 13 for functional analysis. To do this, we utilized *EBA-140* knockout parasites generated in 3D7 (3D7Δ^EBA-140^)^31^. 3D7 is a clonal line derived from NF54^32^, one of the parents in our genetic cross, so provides an appropriate genetic background for interrogating the role of *EBA-140*. We conducted head-to-head competition experiments between 3D7Δ^EBA-140^ and 3D7 wildtype parasites in media containing serum, AlbuMAX or a serum/AlbuMAX mixture to measure fitness consequences of the *EBA-140* knockout. We then used qPCR to quantify frequencies of the two competing parasite lines in each experiment (Fig. 6). 3D7Δ^EBA-140^ showed lower fitness than wildtype parasites in all experiments, but the impact of *EBA-140* deletion on fitness was strongly dependent on media composition. Fitness costs of 3D7^ΔEBA-140^ were low in AlbuMAX (*s* = 0.07±0.04) intermediate in serum: AlbuMAX mixtures (*s* = 0.25±0.02) and high in serum (*s* = 0.61±0.04). Fitness costs resulting from the EBA-140 knockout were significantly higher in serum than in AlbuMAX (Δ*s* = 0.55, *p* = 5.0E-08), indicating that *EBA-140* is the causative gene within the chr.13 QTL.

**Fig 6.**
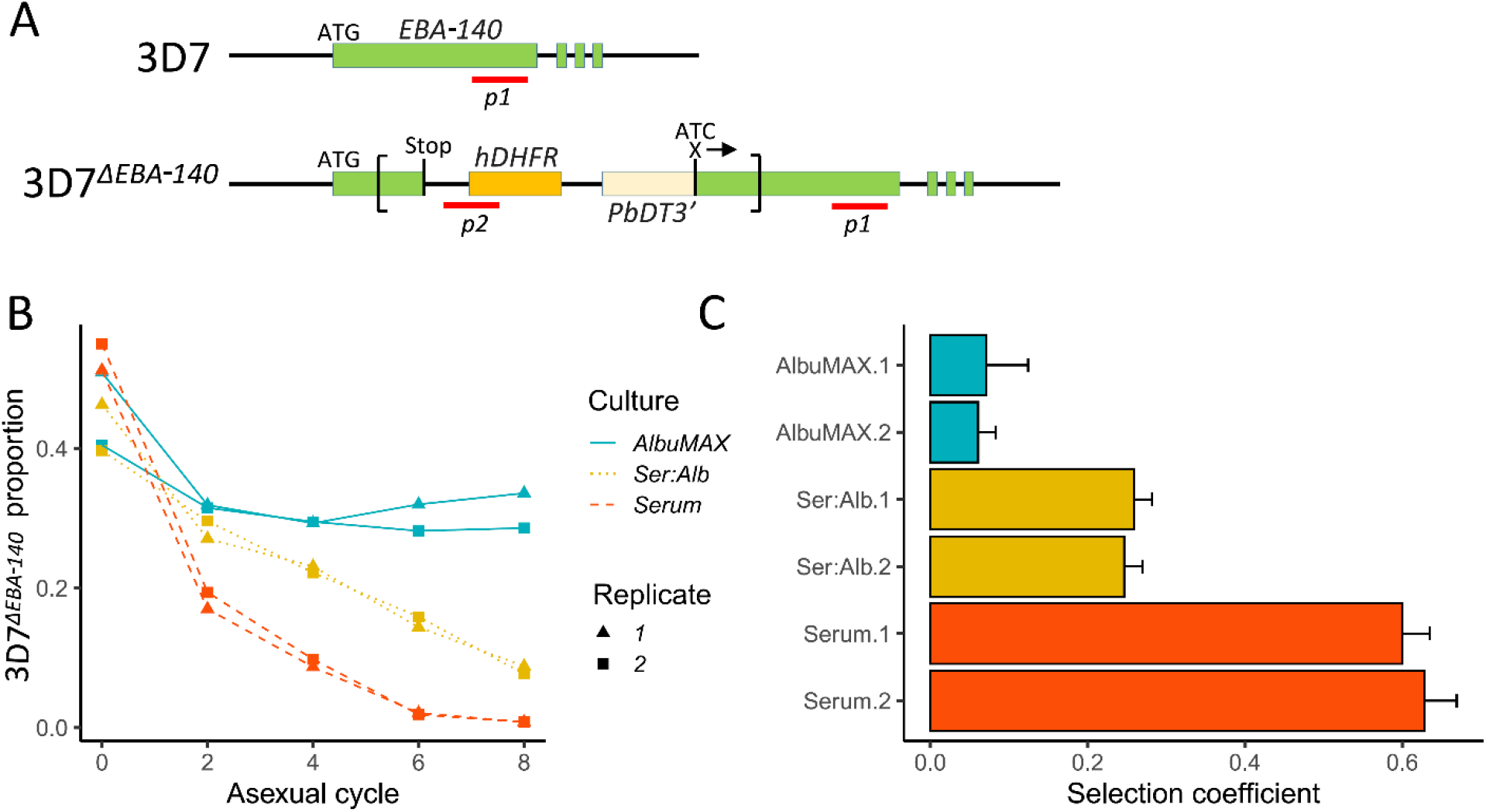
Outcome of competition between 3D7 and 3D7Δ^*EBA-140KO*^ under different culture conditions. (**A**), **Design of qPCR primers**. Top, structure of *EBA-140* gene in *P. falciparum* 3D7 parasite; bottom, disrupted *EBA-140* gene in 3D7 parasite (3D7Δ^*EBA-140*^). The recombinant plasmid pHH1ΔEBA140 (sequence shown inside the square brackets, was integrated into *EBA-140* gene during the generation of 3D7Δ^*EBA-140* 26^. The locations of qPCR amplicons are labeled with red lines. *p1*, located at the 3’ end of *EBA-140*, amplifies both 3D7 and 3D7Δ^*EBA-140*^. *p2* was designed to cover the junction between human dihydrofolate reductase gene (*hDHFR*) (orange box) and a short sequence from the pHH1ΔEBA140 plasmid, which can only be amplified from 3D7Δ^*EBA-140*^. Both *p1* and *p2* primers were designed to not amplify hDHFR from residual human DNA in the culture media. (**B**), Proportion of 3D7Δ^*EBA-140*^ against parasite asexual cycles. (**C**), selection coefficients (*s*) of 3D7Δ^*EBA-140*^ with 1× standard error. Positive values of *s* indicate a disadvantage of 3D7Δ^*EBA-140*^ allele. Culture conditions: Ser:Alb culture media contains both serum and AlbuMAX in a 50:50 ratio.

### Systematic skews observed in both serum and AlbuMAX

We observed three genome regions that showed strong distortions in allele frequency in each independent replicate cross in both human serum and AlbuMAX cultures (Fig. 3), consistent with our earlier report^14, 26^: on chr.7 (named QTL chr.7), in the middle of chr.12 (QTL chr.12) and on the second half of chr. 14 (QTL chr.14.2). We observed strong selection against NHP4026 alleles in the chr.7 and chr.14 QTL regions (*s* = 0.29±0.04 in serum and 0.26±0.04 in AlbuMAX for chr.7, *s* = 0.12±0.02 in both serum and AlbuMAX for chr.14.2), while NF54 alleles were selected against in the chr.12 QTL region, with *s* = 0.30±0.05 in serum and 0.25±0.06 in AlbuMAX (Supplementary Fig. 2). The skews were consistent among all three biological replicates.

The chr.7 QTL spanned from 341 kb to 476 kb (135kb) and contained 33 genes (Supplementary Table 3). The *PfCRT* (PF3D7_0709000), which is known to carry a high fitness cost with CQ resistant alleles^33^, is located at the peak of the chr.7 QTL (Fig 5). Here, NHP4026 carries the CQ resistant *PfCRT* allele while NF54 is CQ sensitive (Supplementary Table 6). In both serum and AlbuMAX cultures, the resistant CQ allele carries an extremely high fitness cost (*s* = 0.29±0.04 in serum and *s* = 0.26±0.04 in AlbuMAX) and is evident as early as day four of *in vitro* culture (Supplementary Fig. 2).

The QTL on chr.12 spanned from 1,141 kb to 1,283 kb (142 kb) and contained 32 genes all expressed in blood stage, of which 9 are high priority candidates. The QTL on chr.14 (numbered 14.2 in Table 1 and Supplementary Table 3) spanned from 2,356 kb to 2,485 kb (129 kb) and contained 23 genes, of which 10 are high priority candidates. These same regions carried high fitness costs in a BSA analysis from an independent genetic cross between two different parental parasites – ART-S MKK2835 and ART-R NHP1337^26^. The multidrug resistance-associated protein 2 (*MRP2*, PF3D7_1229100) and apicoplast ribosomal protein S10 (*ARPS10*, PF3D7_1460900), are both located inside of the chr.12 and chr.14.2 QTL regions respectively. There are total of four non-synonymous mutations and six indels within the *MRP2* locus, and all four parental parasites carry different *MRP2* alleles. *MRP2* alleles from NF54 (from this study) and NHP1337 (from Li et al.^26^) carried high fitness costs during *in vitro* blood cultures with serum containing media. There were two non-synonymous mutations (Val127Met and Asp128His) in the ARPS10 locus (Supplementary Table 4). *ARPS10* alleles with these two mutations (NHP4026 in this study and NHP1337 in previous study by Li et al.^26^) carry a high fitness cost during *in vitro* culture with serum. The Val127Met mutation is one of the genetic background SNPs for *kelch13* alleles on which ART resistance emerged in Cambodia^34^.

## Discussion

### Parasite growth rates and fitness are dependent on growth media

A central finding of this work is that parasite fitness in blood stage culture is strongly dependent on the growth media used. The outcome of competition experiments between the two parental parasites were completely reversed depending on whether media contained serum or AlbuMAX as a lipid source. Surprisingly, the newly isolated parasite, NHP4026, shows high fitness in AlbuMAX, while NF54, which was isolated in 1981^32^ and has a long history of laboratory cultivation using both serum and AlbuMAX based media, shows highest fitness in media containing serum. The fitness traits observed do not appear to be driven by short-term adaptation, and are segregating in the progeny of the genetic crosses, indicative of a genetic basis. The strong association between fitness and growth environment has important implications for experimental malaria biology. Fitness-based assays are widely used for examining impact of piggyBAC insertions^35^, determining the fitness costs of antimalarial drug resistance^36, 37^, and measuring red blood invasion^38^. These results add weight to previous observations that serum or AlbuMAX based media can impact measurements of drug resistance^22^, presentation of antigens on the RBC surface^19^, and highlights the differences between *in vitro* and *in vivo* parasite growth^39, 40^.

### Locus specific selection between serum and AlbuMAX cultures

Because the parental parasites showed distinctive patterns of growth in serum or AlbuMAX, we were able to use genetic crosses, in combination with BSA, to determine the genome regions underlying differential growth in these two media types. We found three QTL regions where allele frequencies of recombinant progeny parasite pools showed dramatic divergence, depending on whether they were grown in serum or in AlbuMAX (Fig. 4). QTLs for differential selection in serum and AlbuMAX were observed in each of the biological replicate recombinant pools and across technical replicates for each pool. Furthermore, selection driving change in allele frequency in these three QTL regions is strong (*s* = 0.10 ± 0.01 [chr.2], *s* = 0.23±0.02 [chr.13] and *s* = 0.12±0.02 [chr.14.1], Table 1).

AlbuMAX is a lipid-loaded bovine serum albumin, while the composition of serum is more complex, containing a variety of proteins and peptides (albumins, globulins, lipoproteins, enzymes and hormones), nutrients (carbohydrates, lipids and amino acids), electrolytes, and small organic molecules. Serum also contains more phospholipid and cholesterol and less fatty acid than AlbuMAX^19^. It has been previously reported that AlbuMAX doesn’t support malaria parasite growth as well as serum^20, 21, 22^. Our analysis reveals three QTLs across the genome that determine parasite growth rate in serum versus AlbuMAX, indicating that this trait has a relatively simple genetic basis in this cross, and suggesting that these QTL regions contain genes involved in nutrient uptake, metabolism, or interactions with serum components. By inspection of the genes under these QTL peaks, we identified three genes – *AST* (QTL chr.2), *EBA-140* (chr.13) and cysteine protease *ATG4* (chr.14.1) – as the strongest candidates (Table 1, full gene listed in Supplementary Table 5).

### Interplay between RBC invasion and growth media

These experiments were initially conceived to identify genes that underlie nutrient utilization/metabolism in *P. falciparum*. However, our functional analysis of the largest QTL (chr.13) reveals that *EBA-140*, a gene known to be involved in RBC invasion^31^, underlies differential growth in media containing human serum or AlbuMAX. EBA-140 mediates *P. falciparum* RBC invasion by binding to the RBC receptor glycophorin C^28^. These results suggest that there are intimate links between the efficiency of RBC invasion and media composition. How might such links be mediated? Multiple studies have showed that mutations in *EBA-140* influence parasite ligand-binding to RBC receptors^28, 30^. In our study, only one amino acid change (Leu112Phe) distinguishes the *EBA-140* from NF54 and NHP4026. We speculate that this mutation influences EBA-140 binding to its host RBC receptor, but that the impact of this mutation is strongly dependent on media-induced changes in the conformation, abundance or accessibility of receptors on RBC surface, or the function of EBA-140 itself during the process of merozoite invasion.

### Candidate genes driving the chr.2 and chr.14.1 QTLs

Some compelling candidate loci that may drive differential growth in serum or AlbuMAX were also found under the two smaller QTLs:

Chr.2: *P. falciparum* acquires nutrients from the host through catabolism of hemoglobin in erythrocytes. During this process, plasmodial AST plays a critical role in the classical tricarboxylic acid (TCA) cycle, and also functions to maintain homeostasis of carbohydrate metabolic pathways^41^. AST is essential^42^, which highlights this gene as a potential bottleneck in energy metabolism and as a target for the design of novel therapeutic strategies. We speculate that selection acts on the regulation of *AST* gene expression levels: we detected no non-synonymous mutations in the coding region, but we found multiple variants in the 5’ UTR and regulatory regions (Supplementary Table 4). However, we cannot exclude the possibility that other neighboring loci may drive the observed allele frequency changes.

Chr.14.1: Cysteine protease ATG4 is essential for autophagy in both yeast, mammals, and protozoan parasites such as *Leishmania major* and *Trypanosoma cruzi* ^43^. Malaria parasites have several cysteine proteases with functions including hemoglobin hydrolysis, RBC rupture, and RBC invasion^44^: two of these (serine repeat antigen 3 and 4) show >2-fold higher expression in AlbuMAX supplemented media^45^. Autophagy involves vesicular trafficking and is important for protein and organelle degradation during cellular differentiation^46^. Conditional knock down of *ATG4* in the apicomplexan *Toxoplasma* leads to growth defects^47^, while autophagy can also be triggered by starvation^46^. In our study, the NF54 *ATG4* allele had higher fitness than NHP4026 during AlbuMAX cultures, while the allele frequencies didn’t change in serum (Supplementary Fig. 2). As AlbuMAX contains fewer components than serum^30^, we speculate that parasites may require higher levels of autophagy to compensate for specific nutrients that are missing in AlbuMAX.

### Replication of fitness-related QTLs in independent genetic crosses

We detected three additional regions of the genome (QTL chr.7, chr.12 and chr.14.2) that show extreme skews in allele frequencies in both serum and AlbuMAX cultures.

The strong chr.7 QTL is almost certainly driven by selection against the chloroquine resistant (CQR) *PfCRT* allele from NHP4026. The same skew was also observed in clones isolated from our previous crosses between the same parental parasites^14^. The rapid change in allele frequency of *PfCRT* alleles equates to selection coefficients (*s*) of 0.29 per asexual life cycle in serum and 0.26 in AlbuMAX (Fig. 5A). These fitness cost estimates are strikingly close to those calculated from laboratory competition experiments that measured fitness costs of the *Dd2*-type *PfCRT* allele to be ~0.3 per asexual life cycle relative to a wild type *PfCRT* alleles ^33^ and are also consistent with decline in frequency of parasites with CQR *PfCRT* observed in Malawi, Kenya and China following the withdrawal of CQ ^33^. Interestingly, NHP4026, the parent carrying the CQR *PfCRT* allele, shows highly competitive fitness in the laboratory, ranking above other SE Asia clinical isolates^23^, and even outcompetes the long-term lab-adapted NF54 bearing the CQ sensitive (CQS) *PfCRT* allele. We speculate that NHP4026 carries compensatory loci that restore fitness in this CQR parasite, but that recombination decouples alleles at *PfCRT* and compensatory loci, resulting in the high fitness cost of *PfCRT* CQR alleles observed in our crosses.

In both our previous genetic cross using different parental parasites^26^ and in the current study, multidrug resistance-associated protein 2 (MRP2) was located at the peak of the chr.12 QTL. *MRP2* belongs to the C-family of ATP binding cassette (ABC) transport proteins that are well known for their role in multidrug resistance. MRP2 mediates the export of drugs, toxins, and endogenous and xenobiotic organic anions^48^, which can thus lead to resistance to multiple drugs. *MRP2* may also contribute to the detoxification of antimalarial drugs. Further experiments are needed to directly determine the function of *MRP2* in parasite fitness during *in vitro* culture.

The chr.14.2 QTL contains the *ARPS10* gene, which is thought to provide a permissive genetic background for emergence of artemisinin resistance^34^ (Supplementary Table 7): this QTL was seen both in this study and our previous cross^26^. Further studies are required to determine whether the mutant *ARPS10* allele might compensate for fitness costs associated with mutant *kelch13* through epistatic interactions.

### Perspectives on bulk segregant approaches for identifying RBC-stage growth-related genes

Our study uses a BSA strategy to systemically identify genes involved in competitive growth, nutrient uptake and metabolism. In contrast, conducting these analyses in a traditional framework, using isolation of individual progeny, measurement of growth phenotypes of each progeny in different media, and genomic characterization of progeny and parents, as required for previous nutritional genetics experiments with *P. falciparum*^7, 49, 50^ is laborious and time consuming. A particular advantage of BSA is that selections are applied to all recombinant progeny simultaneously in a single culture, removing batch effects and experimental variation resulting from conducting parallel measures with individual progeny clones. However, we note that BSA cannot determine the interactions among loci: this can be approached by examining growth of cloned progeny carrying different combinations of parental alleles at the three QTLs.

In this particular experiment, we compared media containing different lipid sources. The chemical differences between serum and AlbuMAX are complex, so it will be difficult to identify the precise media components driving differential selection. However, following the example of bacterial genetics where selective media has played a central role ^51, 52, 53^, we envisage that screening of parental parasites in media that differ in levels of single nutrients will identify parasites with differing nutrient requirements. Subsequent genetic crosses and BSA experiments can then localize the genes underlying these nutritional polymorphisms. Such experiments can be further extended to examine RBC invasion phenotypes by conducting BSA experiments using RBCs carrying different invasion receptors: such RBCs can now be efficiently generated using CRISPR/Cas9 editing of hemopoietic stem cells^54^, so other aspects of host genetics can be maintained constant. We conclude that genetic crosses and BSA genetic cross can provide a versatile approach for dissecting the genetic basis of nutrient acquisition, and RBC invasion determining parasite growth, and for examining host parasite interactions at the molecular level.

## Methods

### Ethics approval and consent to participate

The study was performed in strict accordance with the recommendations in the Guide for the Care and Use of Laboratory Animals of the National Institutes of Health (NIH), USA. To this end, the Seattle Children’s Research Institute (SCRI) has an Assurance from the Public Health Service (PHS) through the Office of Laboratory Animal Welfare (OLAW) for work approved by its Institutional Animal Care and Use Committee (IACUC). All of the work carried out in this study was specifically reviewed and approved by the SCRI IACUC.

### Culture media with serum and AlbuMAX

Contents and manufacturers of the culture media used in this study for *P. falciparum* are as listed in Supplementary Table 8. In summary, we used RPMI 1640 as basal medium. We added 2 mM L-glutamine as amino acid supplement, 25mM HEPES for maintaining culture pH and 50 μM hypoxanthine as a nutrient additive which helps in cell growth. To prevent the growth of fungi and bacteria, we used 50 IU/ml penicillin, 50 μg/ml streptomycin and 50 μg/ml vancomycin for the first two asexual life cycles (4 days). The basal medium was supplemented either with 10% O+ human serum or with 0.5 % AlbuMAX II. O+ erythrocytes were added every two days into the culture media to support amplification of the parasite population. We maintained all the cultures at 37°C with 5% O2, 5% CO2, and 90% N2, with 2% hematocrit. Only one batch of reagents were used through the whole experiment.

### Preparation of the genetic cross

We generated the cross using FRG NOD huHep mice with human chimeric livers and *A. stephensi* mosquitoes as described in Vaughan *et al* ^13^ and Li et al ^26^ (Fig 2A). We used NF54 (lab adapted Africa parasite) and NHP4026 (newly cloned clinical isolate from the Thai-Myanmar border) in this study. Gametocytes from both the parasite strains were diluted to 0.5% gametocytemia using human serum erythrocyte mix, to generate infectious blood meals (IBMs). IBMs from each parent was mixed at equal ratio and fed to ~450 mosquitos (3 cages of 150 mosquitoes, of which ~45 were mosquitoes were sacrificed for prevalence test). Recombinants are generated after gametes fuse to form zygotes in the mosquito midgut (Fig. 2A). Replication of the four meiotic products ultimately leads to the generation of thousands of haploid sporozoites within each oocyst. We examined the mosquito infection rate and oocyst number per infected mosquito 7 days post-feeding. Fifteen mosquitoes were randomly picked from each cage and dissected under microscopy. For this cross, the oocyst prevalence was 93% (range: 86-100%), with an average burden of 14 oocysts per mosquito midgut (range: 2-61). To ensure that recombinants from each pool were independent, we injected pooled sporozoites into each of three FRG huHep mice from a different cage of mosquitoes (~100). We generated three independent recombinant pools from this experiment.

Sporozoites are isolated from infected mosquito salivary glands and 2-4 Million sporozoites from each cage of mosquitoes were injected into three FRG huHep mice (one cage per mouse), intravenously. To allow the liver stage-to-blood stage transition, mice are infused with human erythrocytes six and seven days after sporozoite injection. Four hours after the second infusion, the mice are euthanized and exsanguinated to isolate the circulating ring stage *P. falciparum*-infected human erythrocytes. The parasites from each mouse constitute the initial recombinant pools for further segregation experiment. All initial recombinant pools were maintained using AlbuMAX culture for 24hr to stabilize the newly transitioned ring stage parasites. We prepared three recombinant pools in this study.

The expected proportion of oocysts that generated from mating between male and female gametes of the same genotype (selfed) is 0.5. The allele frequencies of NF54 in recombinant pools of parasites emerging from the liver were 0.52±0.001, 0.50±0.002 and 0.55±0.002 for the three mice. Combining all the information, we estimate that there were 28 (14 oocysts × 4 recombinants per oocyst × 0.5 selfed) recombinant genotypes per infected mosquito, which gave us ~2800 (28 × ~100 mosquitoes) unique recombinants per pool.

### Sample collection and sequencing

We aliquoted each initial recombinant pool into four cultures and maintained two of them with serum medium and the other two with AlbuMAX medium (Fig 2B and Supplementary Table 8). There were total of 12 cultures (3 mice as biological replicates × [serum + AlbuMAX II] × 2 technical replicates). We maintained all the cultures with standard six-well plate for 34 days. Freshly packed erythrocytes were added every 2 days to each replicate, meanwhile parasites cultures were diluted to 1% parasitemia to avoid stressing. 70ul packed erythrocytes were collected and frozen down every 2-4 days. We extracted and purified genomic DNA using the Qiagen DNA mini kit, and quantified amounts using Qubit. We constructed next generation sequencing libraries using 50-100 ng DNA or sWGA product following the KAPA HyperPlus Kit protocol with 3-cycle of PCR. All libraries were sequenced at 150bp pair-end to a minimum coverage of 100× using Illumina Novaseq S4 or Hiseq X sequencers.

### Genotype calling

We genotyped the parental strains and bulk populations as described in ^26^. We first generated a “mock” genome according to the genotype of parent NF54. We mapped the whole-genome sequencing reads both from parental parasites and progeny against this mock genome using BWA mem (http://bio-bwa.sourceforge.net/) under the default parameters. We excluded the high variable genome regions (subtelomeric repeats, hypervariable regions and centromeres) and only performed genotype calling in the 21 Mb core genome (defined in ^25^). The resulting alignments were then converted to SAM format, sorted to BAM format, and deduplicated using picard tools v2.0.1 (http://broadinstitute.github.io/picard/). We used Genome Analysis Toolkit GATK v3.7 (https://software.broadinstitute.org/gatk/) to recalibrate the base quality score based on a set of verified known variants ^25^. We called variants using HaplotypeCaller and then merged using GenotypeGVCFs with default parameters except for sample ploidy 1.

We only applied filters to the GATK genotypes of parental parasites, using standard filter methods described by McDew-White et al ^55^. The recalibrated variant quality scores (VQSR) were calculated by comparing the raw variant distribution with the known and verified *Plasmodium* variant dataset, and loci with VQSR less than 1 were removed from further analysis. After filtration, we selected SNP loci that are distinct in two parents, and only used those for further bulk segregant analysis.

### Bulk segregant analysis

We performed the BSA as described in^26^. Only loci with coverage > 30× were used for bulk segregant analysis. We counted reads with genotypes of each parent and calculated allele frequencies at each variable locus. Allele frequencies of NF54 were plotted across the genome, and outliers were removed following Hampel’s rule ^56^ with a window size of 100 loci. We performed the BSA analyses using the R package QTLseqr ^57^. Extreme-QTLs were defined as regions with G’ > 20 ^58^. Once a QTL was detected, we calculated and approximate 95% confidence interval using Li’s method ^59^ to localize causative genes. We also measured the fitness cost at each mutation by fitting a linear model between the natural log of the allele ratio (freq[allele1]/freq[allele2]) against time (measured in 48hr parasite asexual cycles). The slope provides a measure of the selection coefficient (*s*) driving each mutation ^60^. The raw *s* values were tricube-smoothed with a window size of 100 kb to remove noise ^61, 62^.

### Head-to-head competition experiments

We conducted head-to-head competition experiments between parental NF54 and NHP4026 parasites, and between the 3D7Δ^*EBA-140*^ and the 3D7 wildtype parasites. For each comparison, we synchronized the parasites at ring stages. These parasites were adapted to respective media formulations for 2 asexual replication cycles. Two competitor parasites were then mixed in 1:1 ratio at ring stages with final 1% parasitemia and cultured as two technical replicates in respective media formulation. We maintained two of the aliquots in human serum supplemented media, two in AlbuMAX supplemented media, and two in medium containing equal amount of serum and AlbuMAX. All competition assays were conducted in 6-well plates, with 5 ml culture volume. All cultures received media changes every day and cultures were cut to 1% parasitemia every other replication cycle. Samples were collected every 4 days to monitor the outcome of each competition and frozen at −80 °C until genomic DNA was extracted.

To determine the frequency of each parental parasite (NF54 or NHP4026) competition, we used a microsatellite-based method developed by Tirrell et al^23^. PCR amplification products from microsatellite TA119 (forward primer: TCCTCGATTATATTATTGCA; reverse primer: TAATACATTCCCATTAGATC) were analyzed using Applied Biosystems 3730xI DNA Analyzer (ThermoFisher). Relative density of amplicons from each parent was then scored by the height of corresponding fluorescent peak comparing to the overall signal.

For 3D7Δ^*EBA-140*^ and wildtype parasites competition, we used a qPCR assay to determine the frequency of each parasite. The 3D7Δ^*EBA-140*^ parasites were generated by Maier et al., using transfection plasmid pHH1ΔEBA140 containing human dihydrofolate reductase gene (*hDHFR*) sequence^31^. The plasmid sequences were integrated into the genome of 3D7Δ^*EBA-140*^ parasites and are absent in wildtype parasites. We designed qPCR primers (*p1*-forward: CGGAATGGCGAATAACAATG; *p1*-reverse: ATTCCCCGCACTTTCCTCTT) to amplify genomic regions for both 3D7Δ^*EBA-140*^ and wildtype parasites; we used another set of primers (*p2*-forward: TGATGTCCAGGAGGAGAAAGG; *p2*-reverse: TCGCTATCCCATAAATTACAAAACA) which cover the junction between *hDHFR* and a short sequence from the pHH1ΔEBA140 plasmid, to amplify genome regions only from 3D7Δ^*EBA-140*^ parasites (Fig. 6A). Both *p1* and *p2* primers were not able to amplify using human DNA residual as template. The 3D7Δ^*EBA-140*^ allele frequency was then calculated as copies of 3D7Δ^*EBA-140*^ parasite genomes/copies of total parasite genomes.

## Data Availability

All data needed to evaluate the conclusions in the paper are present in the paper and/or the Supplementary Materials. All raw sequencing data have been submitted to the NABI Sequence Read Archive (SRA, https://www.ncbi.nlm.nih.gov/sra) under the project number of PRJNA524855. Additional data related to this paper may be requested from the authors.

## Acknowledgments

This work was supported by National Institutes of Health (NIH) program project grant P01 AI127338 (to MF), by NIH grant R37 AI048071 (to TJCA) and NIH grant R21 AI133369 (to AMV). Work at Texas Biomedical Research Institute was conducted in facilities constructed with support from Research Facilities Improvement Program grant C06 RR013556 from the National Center for Research Resources. SMRU is part of the Mahidol Oxford University Research Unit supported by the Wellcome Trust of Great Britain.

## Author contributions

S.K., A.M.V., X.L. and T.J.C.A. designed the experiments. S.K., M.T.H, B.A.A., S.Y.K and N.C. prepared the crosses and collected samples. M.M.W, A.R., E.D. and A.S. prepared the genomic DNA libraries. F.N. provided the parental parasite from Thailand. L.A.C cloned the parental parasites. X.L. performed all the NGS analysis and data curation. S.K., X.L. and T.J.C.A. wrote the original manuscript. K.V.B, K.A.B, I.H.C, S.H.I.K, F.N., M.T.F and A.M.V reviewed and edited the manuscript.

## Competing interests

The authors declare that they have no competing interests.

## Supporting information

**Supplementary Figure 1. Selection coefficients (*s*) across the genome**. Estimation of *s* was based on the changes of allele frequency from day1 to day30 of cultures. Positive values of *s* indicate a disadvantage for alleles inherited from NHP4026. Red and green lines indicate cultures by serum and AlbuMAX.

**Supplementary Fig. 2. Estimation of selection coefficients from the changes in allele frequencies in candidate gene regions**. **(A-F)**, NF54 allele frequency for gene regions of *CRT*, *MRP2*, *ARPS10*, *AST*, *EBA-10* and *ATG*, separately. Selection coefficients (*s*) were calculated as the slope of the linear model between the natural log of the allele ratio [freq (NF54)/freq (NHP4026)] against time. Positive values of *s* indicate a disadvantage for alleles inherited from NHP4026.

**Supplementary Figure 3. NF54 allele frequency at candidate gene regions in world-wide malaria parasite populations**. (A-C), single nucleotide polymorphisms for gene *AST*, *EBA-10* and *ATG4*, separately. WAF: west Africa, EAF: east Africa, CAF: central Africa, SAM: south America, ESEA: east Southeast (SE) Asia, SAS: south Asia, WSEA: west SE Asia, OCE: Pacific Ocean. The allele frequency analysis was performed using genomic database for *Plasmodium falciparum* (MalariaGEN, https://www.malariagen.net/).

**Supplementary Table 1**. SNPs between NF54 and NHP4026 parental parasites.

**Supplementary Table 2**. Summary of cloned progeny from cross between NF54 and NHP4026.

**Supplementary Table 3**. Genes inside the QTL regions.

**Supplementary Table 4**. SNPs and indels inside of the QTL regions.

**Supplementary Table 5**. Top candidate genes located inside of the QTL regions. A full list of genes found within QTL regions is shown in Supplementary Table 3.

**Supplementary Table 6**. Summary of selection coefficients (*s*) at candidate gene regions.

**Supplementary Table 7**. Genotypes of parental parasites.

**Supplementary Table 8**. Summary of media components used in this study.

## Notes

### Competing Interest Statement

The authors have declared no competing interest.

### Summary of Updates

Title has been changed New results added: Figure 1 now shows competitive growth phenotypes of the two parents used in the genetic crosses described. Figure 6 now shows validation experiments for one of the QTLs identified. Extensive changes to the results and discussion reflect new results obtained

## References

1. Draper SJ, et al. Recent advances in recombinant protein-based malaria vaccines. Vaccine 33, 7433–7443 (2015).

2. Birnbaum J, et al. A Kelch13-defined endocytosis pathway mediates artemisinin resistance in malaria parasites. Science 367, 51–59 (2020).

3. Wang J, et al. Haem-activated promiscuous targeting of artemisinin in Plasmodium falciparum. Nat Commun 6, 10111 (2015).

4. Yang J, et al. Advances in the research on the targets of anti-malaria actions of artemisinin. Pharmacol Ther 216, 107697 (2020).

5. Shafik SH, et al. The natural function of the malaria parasite’s chloroquine resistance transporter. Nat Commun 11, 3922 (2020).

6. Gregson A, Plowe CV. Mechanisms of resistance of malaria parasites to antifolates. Pharmacol Rev 57, 117–145 (2005).

7. Nguitragool W, et al. Malaria parasite clag3 genes determine channel-mediated nutrient uptake by infected red blood cells. Cell 145, 665–677 (2011).

8. Wang P, Brobey RK, Horii T, Sims PF, Hyde JE. Utilization of exogenous folate in the human malaria parasite Plasmodium falciparum and its critical role in antifolate drug synergy. Mol Microbiol 32, 1254–1262 (1999).

9. Ehrenreich IM, et al. Dissection of genetically complex traits with extremely large pools of yeast segregants. Nature 464, 1039–1042 (2010).

10. Pattaradilokrat S, Culleton RL, Cheesman SJ, Carter R. Gene encoding erythrocyte binding ligand linked to blood stage multiplication rate phenotype in Plasmodium yoelii yoelii. Proc Natl Acad Sci U S A 106, 7161–7166 (2009).

11. Abkallo HM, et al. Rapid identification of genes controlling virulence and immunity in malaria parasites. PLoS Pathog 13, e1006447 (2017).

12. Hunt P, et al. Experimental evolution, genetic analysis and genome re-sequencing reveal the mutation conferring artemisinin resistance in an isogenic lineage of malaria parasites. BMC Genomics 11, 499 (2010).

13. Vaughan AM, et al. Plasmodium falciparum genetic crosses in a humanized mouse model. Nat Methods 12, 631–633 (2015).

14. Button-Simons KA, et al. The power and promise of genetic mapping from Plasmodium falciparum crosses utilizing human liver-chimeric mice. Commun Biol 4, 734 (2021).

15. Vendrely KM, Kumar S, Li X, Vaughan AM. Humanized Mice and the Rebirth of Malaria Genetic Crosses. Trends Parasitol 36, 850–863 (2020).

16. Trager W, Jensen JB. Human malaria parasites in continuous culture. Science 193, 673–675 (1976).

17. Moore GE, Gerner RE, Franklin HA. Culture of normal human leukocytes. JAMA 199, 519–524 (1967).

18. Sherman IW. Transport of amino acids and nucleic acid precursors in malarial parasites. Bull World Health Organ 55, 211–225 (1977).

19. Frankland S, et al. Serum lipoproteins promote efficient presentation of the malaria virulence protein PfEMP1 at the erythrocyte surface. Eukaryot Cell 6, 1584–1594 (2007).

20. Basco LK. Molecular epidemiology of malaria in cameroon. XX. Experimental studies on various factors of in vitro drug sensitivity assays using fresh isolates of Plasmodium falciparum. Am J Trop Med Hyg 70, 474–480 (2004).

21. Dohutia C, Mohapatra PK, Bhattacharyya DR, Gogoi K, Bora K, Goswami BK. In vitro adaptability of Plasmodium falciparum to different fresh serum alternatives. J Parasit Dis 41, 371–374 (2017).

22. Ringwald P, Meche FS, Bickii J, Basco LK. In vitro culture and drug sensitivity assay of Plasmodium falciparum with nonserum substitute and acute-phase sera. J Clin Microbiol 37, 700–705 (1999).

23. Tirrell AR, et al. Pairwise growth competitions identify relative fitness relationships among artemisinin resistant Plasmodium falciparum field isolates. Malar J 18, 295 (2019).

24. Preston MD, et al. A barcode of organellar genome polymorphisms identifies the geographic origin of Plasmodium falciparum strains. Nat Commun 5, 4052 (2014).

25. Miles A, et al. Indels, structural variation, and recombination drive genomic diversity in Plasmodium falciparum. Genome research 26, 1288–1299 (2016).

26. Li X, et al. Genetic mapping of fitness determinants across the malaria parasite Plasmodium falciparum life cycle. PLoS Genet 15, e1008453 (2019).

27. Toney MD. Aspartate aminotransferase: an old dog teaches new tricks. Arch Biochem Biophys 544, 119–127 (2014).

28. Adams JH, Sim BK, Dolan SA, Fang X, Kaslow DC, Miller LH. A family of erythrocyte binding proteins of malaria parasites. Proc Natl Acad Sci U S A 89, 7085–7089 (1992).

29. Gomez-Escobar N, Amambua-Ngwa A, Walther M, Okebe J, Ebonyi A, Conway DJ. Erythrocyte invasion and merozoite ligand gene expression in severe and mild Plasmodium falciparum malaria. J Infect Dis 201, 444–452 (2010).

30. Lin DH, Malpede BM, Batchelor JD, Tolia NH. Crystal and solution structures of Plasmodium falciparum erythrocyte-binding antigen 140 reveal determinants of receptor specificity during erythrocyte invasion. J Biol Chem 287, 36830–36836 (2012).

31. Maier AG, et al. Plasmodium falciparum erythrocyte invasion through glycophorin C and selection for Gerbich negativity in human populations. Nat Med 9, 87–92 (2003).

32. Walliker D, et al. Genetic analysis of the human malaria parasite Plasmodium falciparum. Science 236, 1661–1666 (1987).

33. Ecker A, Lehane AM, Clain J, Fidock DA. PfCRT and its role in antimalarial drug resistance. Trends Parasitol 28, 504–514 (2012).

34. Miotto O, et al. Genetic architecture of artemisinin-resistant Plasmodium falciparum. Nat Genet 47, 226–234 (2015).

35. Zhang M, et al. Uncovering the essential genes of the human malaria parasite Plasmodium falciparum by saturation mutagenesis. Science 360, (2018).

36. Stokes BH, et al. Plasmodium falciparum K13 mutations in Africa and Asia impact artemisinin resistance and parasite fitness. Elife 10, (2021).

37. Nair S, et al. Fitness Costs and the Rapid Spread of kelch13-C580Y Substitutions Conferring Artemisinin Resistance. Antimicrob Agents Chemother 62, (2018).

38. Ebel ER, Kuypers FA, Lin C, Petrov DA, Egan ES. Common host variation drives malaria parasite fitness in healthy human red cells. Elife 10, (2021).

39. Brown AC, Guler JL. From Circulation to Cultivation: Plasmodium In Vivo versus In Vitro. Trends Parasitol 36, 914–926 (2020).

40. LeRoux M, Lakshmanan V, Daily JP. Plasmodium falciparum biology: analysis of in vitro versus in vivo growth conditions. Trends Parasitol 25, 474–481 (2009).

41. Wrenger C, Muller IB, Silber AM, Jordanova R, Lamzin VS, Groves MR. Aspartate aminotransferase: bridging carbohydrate and energy metabolism in Plasmodium falciparum. Curr Drug Metab 13, 332–336 (2012).

42. Ting LM, et al. Targeting a novel Plasmodium falciparum purine recycling pathway with specific immucillins. J Biol Chem 280, 9547–9554 (2005).

43. Siqueira-Neto JL, et al. Cysteine proteases in protozoan parasites. PLoS Negl Trop Dis 12, e0006512 (2018).

44. Rosenthal PJ. Cysteine proteases of malaria parasites. Int J Parasitol 34, 1489–1499 (2004).

45. Singh K, Agarwal A, Khan SI, Walker LA, Tekwani BL. Growth, drug susceptibility, and gene expression profiling of Plasmodium falciparum cultured in medium supplemented with human serum or lipid-rich bovine serum albumin [corrected]. J Biomol Screen 12, 1109–1114 (2007).

46. Levine B, Klionsky DJ. Development by self-digestion: molecular mechanisms and biological functions of autophagy. Dev Cell 6, 463–477 (2004).

47. Kong-Hap MA, et al. Regulation of ATG8 membrane association by ATG4 in the parasitic protist Toxoplasma gondii. Autophagy 9, 1334–1348 (2013).

48. Nies AT, Keppler D. The apical conjugate efflux pump ABCC2 (MRP2). Pflugers Arch 453, 643–659 (2007).

49. Gupta A, et al. Complex nutrient channel phenotypes despite Mendelian inheritance in a Plasmodium falciparum genetic cross. PLoS Pathog 16, e1008363 (2020).

50. Gupta A, Balabaskaran-Nina P, Nguitragool W, Saggu GS, Schureck MA, Desai SA. CLAG3 Self-Associates in Malaria Parasites and Quantitatively Determines Nutrient Uptake Channels at the Host Membrane. mBio 9, (2018).

51. Maheshwari R. Joshua Lederberg (1925–2008) and the pink–orange fungus Neurospora. Current Science 101, 687–689 (2011).

52. Shuman HA, Silhavy TJ. The art and design of genetic screens: Escherichia coli. Nat Rev Genet 4, 419–431 (2003).

53. Ryan FJ. Back-mutation and adaptation of nutritional mutants. In: Cold Spring Harbor Symposia on Quantitative Biology). Cold Spring Harbor Laboratory Press (1946).

54. Scully EJ, et al. Generation of an immortalized erythroid progenitor cell line from peripheral blood: A model system for the functional analysis of Plasmodium spp. invasion. Am J Hematol 94, 963–974 (2019).

55. McDew-White M, Li X, Nkhoma SC, Nair S, Cheeseman I, Anderson TJ. Mode and tempo of microsatellite length change in a malaria parasite mutation accumulation experiment. Genome Biology and Evolution 11, 1971–1985 (2019).

56. Davies L, Gather U. The identification of multiple outliers. Journal of the American Statistical Association 88, 782–792 (1993).

57. Mansfeld BN, Grumet R. QTLseqr: An R package for bulk segregant analysis with next-generation sequencing. The Plant Genome 11, 180006 (2018).

58. Magwene PM, Willis JH, Kelly JK. The statistics of bulk segregant analysis using next generation sequencing. PLoS computational biology 7, e1002255 (2011).

59. Li H. A quick method to calculate QTL confidence interval. Journal of genetics 90, 355–360 (2011).

60. Dykhuizen D, Hartl DL. Selective neutrality of 6PGD allozymes in E. coli and the effects of genetic background. Genetics 96, 801–817 (1980).

61. Nadaraya EA. On estimating regression. Theory of Probability & Its Applications 9, 141–142 (1964).

62. Watson GS. Smooth regression analysis. Sankhyā: The Indian Journal of Statistics, Series A, 359–372 (1964).

